# Nephrotic Syndrome-Associated Hypercoagulopathy is Alleviated by Nuclear Receptor Agonist Therapy with both Pioglitazone and Glucocorticoids

**DOI:** 10.1101/2020.01.07.897751

**Authors:** Amanda P. Waller, Shipra Agrawal, Katelyn J. Wolfgang, Jiro Kino, Melinda A. Chanley, William E. Smoyer, Bryce A. Kerlin, the Pediatric Nephrology Research Consortium (PNRC)

## Abstract

**Background:** Thrombosis is a potentially life-threatening nephrotic syndrome (NS) complication. We have previously demonstrated that hypercoagulopathy is proportional to NS severity in rat models and that pioglitazone (Pio) reduces proteinuria both independently and in combination with methylprednisolone (MP), a glucocorticoid (GC). However, the effect of these treatments on NS-associated hypercoagulopathy remains unknown. We thus sought to determine the ability of Pio and GC to alleviate NS-associated hypercoagulopathy.

**Methods:** Puromycin aminonucleoside-induced rat NS was treated with sham, Low- or High-dose MP, Pio, or combination (Pio+Low-MP) and plasma was collected at day 11. Plasma samples were collected from children with steroid-sensitive NS (SSNS) and steroid-resistant NS (SRNS) upon presentation and after 7 weeks of GC therapy. Plasma endogenous thrombin potential (ETP), antithrombin (AT) activity, and albumin (Alb) were measured using thrombin generation, amidolytic, and colorimetric assays, respectively.

**Results:** In a rat model of NS, both High-MP and Pio improved proteinuria and corrected hypoalbuminemia, ETP and AT activity (*P*<0.05). Proteinuria (*P*=0.005) and hypoalbuminemia (*P*<0.001) were correlated with ETP. In childhood NS, while ETP was not different at presentation, GC therapy improved proteinuria, hypoalbuminemia, and ETP in children with SSNS (*P*<0.001) but not SRNS (*P*=0.330).

**Conclusions:** Both Pio and GC diminish proteinuria and significantly alleviate hypercoagulopathy. Both Pio and MP improved hypercoagulopathy in rats, and successful GC therapy (SSNS) also improved hypercoagulopathy in childhood NS. These data suggest that even a partial reduction in proteinuria may reduce NS-associated thrombotic risk.

**SIGNIFICANCE STATEMENT:** Nephrotic syndrome (NS) is characterized by massive proteinuria and is complicated by a complex, acquired hypercoagulopathy that markedly increases the risk for potentially life-threatening venous thromboembolism (VTE). This study demonstrates a strong correlation between proteinuria reduction and improvement of an established VTE-risk biomarker, in both a well-established animal model and in childhood NS before and after steroid treatment. We show that nuclear receptor agonists with known disparate mechanisms of action successfully reduce proteinuria and simultaneously improve NS-associated hypercoagulopathy. These data suggest that complete or partial proteinuria reduction by any therapeutic modality may concurrently reduce NS-associated thrombotic risk.

## INTRODUCTION

Nephrotic syndrome (NS) is characterized by glomerular injury and massive urinary protein loss, which leads to a severe, acquired hypercoagulopathy associated with an elevated risk for life-threatening venous thromboembolic (VTE) disease that afflicts up to 25% of NS patients.^1–4^ Epidemiologic studies have demonstrated that both proteinuria severity and hypoalbuminemia are independently predictive of NS-related VTE-risk.^1,5–7^ However, the indications for anticoagulant prophylaxis in the setting of NS remain ill-defined and controversial.^8–11^ Moreover, there is no consensus on when it is safe to discontinue anticoagulation. We have previously demonstrated, in animal models of NS, that disease severity (proteinuria and hypoalbuminemia) is tightly correlated with endogenous thrombin potential (ETP), a global measure of hypercoagulopathy that is predictive of VTE-risk, but not yet widely available for clinical use.^12–23^ Therefore, proteinuria and serum albumin levels, which are routinely followed biomarkers of NS disease activity, may become useful surrogate markers of VTE-risk and thus guide clinical trials of anticoagulant prophylaxis in NS.

Meanwhile, the extent to which disease treatment alters NS-hypercoagulopathy and, ultimately, modulation of VTE-risk remains unknown. Current therapeutic options for NS include glucocorticoids (e.g. methylprednisolone; MP) and other immunosuppressive agents, which are associated with significant toxicity and have limited efficacy.^24–28^ There is thus a clear need to develop novel NS therapeutics.^25, 29^ Glucocorticoids (GC) are thought to modulate their effects primarily via activation of glucocorticoid receptor, which is a member of the nuclear receptor superfamily. We and others have thus investigated thiazolidinediones, which are agonists of an alternative nuclear receptor (peroxisome proliferator-activated receptor gamma [PPARγ]) for treatment of NS.^30–32^ It has previously been demonstrated that pioglitazone (Pio), a thiazolidinedione that is FDA-approved for treatment of type 2 diabetes mellitus, also reduces proteinuria in rat models of NS (both independently and in combination with MP).^30, 32, 33^ Some studies suggest that the successful induction of remission may improve or normalize various aspects of the complex hemostatic derangements observed in human NS.^1^ Meanwhile, both MP and Pio may have NS-independent effects on the hemostatic system.^34–42^ Thus, the ultimate net effects of MP or Pio on NS-associated hypercoagulopathy remain poorly defined.

Based upon these observations, we hypothesized that efficacious NS therapies that act through nuclear receptor signaling would simultaneously reduce proteinuria, improve hypoalbuminemia, and alleviate hypercoagulopathy. If so, these data may begin to inform the relationship between NS disease activity and VTE-risk which, in turn, may guide more appropriate and judicious use of anticoagulant prophylaxis in patients with NS. To test this hypothesis, we designed experiments to determine the effects of Pio and MP on ETP, both in rats with puromycin aminonucleoside (PAN) induced nephrosis and in healthy rats. We further explored this hypothesis by determining the effects of GC on ETP before and after treatment in a cohort of children with newly-diagnosed NS. Here we show that nuclear receptor agonist therapies that effectively reduce NS-associated proteinuria and hypoalbuminemia simultaneously reduce hypercoagulopathy in both PAN-induced rat NS and childhood NS.

## METHODS

### Puromycin Aminonucleoside Rat Nephrosis

The data presented in this report, were derived from both banked samples from our previously reported experiments and additional healthy and PAN-NS rats.^30^ In our previous report, plasma samples for coagulation assays were not collected from all animals. Thus, the data presented herein represent analyses of the animals with available plasma, supplemented with additional animals as required to adequately power the coagulation studies. All procedures were approved by the Institutional Animal Care and Use Committee, in accordance with the NIH Guide for the Care and Use of Laboratory Animals. A summary of the *in vivo* experiments and their adherence to the ARRIVE Guidelines is provided in Table S1. Male Wistar rats (body weight ~150 g, age ~45-50 d) received a single tail vein injection of PAN (50 mg/kg; 5 groups) or saline (4 groups) on “Day 0.” PAN-induced proteinuria was treated daily with sham, Low- or High-dose MP (methylprednisolone; 5 or 15 mg/kg via intraperitoneal injection; hereafter, “Low-MP” or “High-MP”), Pio (10 mg/kg via oral gavage), or combination therapy with Pio and Low-MP (hereafter, “Pio+Low-MP”), as previously reported, followed by euthanasia 24-28 hours after the final dose (*n*=8-13/group).^30^ In order to further confirm the sensitivity of PAN-NS hypercoagulopathy to High-MP, we also investigated a range of proteinuria levels obtained by varying the PAN dose and administration route.^12, 43^ For these experiments we used six groups (*n*=4-6/ group) of male Wistar rats (weight ~150-200 g) which received a single dose of saline (*n*=8 controls) or PAN, 75 mg/kg via either tail vein (IV) or intraperitoneal (IP) injection, or 100 mg/kg IP. We also compared the effects of High-MP, Pio, and Pio+Low-MP treatment in a set of healthy rats not given PAN (*n*=4/group). Morning spot urine samples were collected on Days 0 (before PAN injection) and 11 for urinary protein:creatinine ratio (UPC) analysis. On day 11, the rats were anesthetized with 3% isoflurane and blood was collected from the inferior vena cava through a 23-G needle into a final concentration of 0.32% NaCitrate/1.45 μM Corn Trypsin Inhibitor (CTI; Haematologic Technologies Inc., Vermont, VT, USA), processed to Platelet Poor Plasma (PPP) as previously described, and stored at −80°C until analyzed.^12^ UPC was measured by Antech Diagnostics (Morrisville, NC), using standard techniques that are fully compliant with Good Laboratory Practice regulations.^12^ Plasma albumin concentrations were determined using a bromocresol purple (BCP) assay (QuantiChrom BCP; BioAssay Systems, Hayward, CA).

### Pediatric Nephrology Research Consortium Cohort

Children with incident NS were recruited from Pediatric Nephrology Research Consortium (PNRC) participating centers (see participating PNRC centers and investigators in Appendix). The study protocols and consent documents were approved by the Nationwide Children’s Hospital Institutional Review Board (IRB05-00544, IRB07-004, & IRB12-00039) and at each participating PNRC center. Glucocorticoid (GC) therapy naïve children 1 − 18 years of age presenting with edema and proteinuria ≥3+ by dipstick were eligible for enrolment. Steroid sensitive NS (SSNS) was defined as disease remission following 7 (± 0.4) weeks of standard-of-care GC therapy (dose and formulation at the treating physician’s discretion), whereas steroid resistant NS (SRNS) was defined as failure to achieve complete remission during this time frame, as determined by resolution of proteinuria by urine dipstick or UPC. Blood was collected at the time of enrolment (prior to GC exposure) and a paired blood specimen was obtained after 7 (± 0.4) weeks of GC therapy at the time steroid-responsiveness was assessed. Importantly, all of the children were still on GC therapy when the second sample was obtained. Blood was collected into 0.1 M sodium citrate cell preparation tubes containing a cell separator system (BD Vacutainer CPT (REF 362761) Becton, Dickinson and Company, Franklin Lakes, NJ) at an 8:1 blood-to-citrate ratio (~0.33% final concentration NaCitrate). After processing, plasma was frozen at −80°C and transferred to the Abigail Wexner Research Institute at Nationwide Children’s for analysis. Select demographics from this pediatric cohort at disease presentation are provided in Table S2.

### Coagulation Parameters

ETP was determined on PPP (neat and diluted 1:1 with buffer, for patients and rats, respectively) using the Technothrombin TGA kit (Technoclone, Vienna, Austria) and TGA RC low reagent, and read on a Spectramax M2 fluorescent plate reader (Molecular Devices, Sunnyvale, California), as previously described.^12^ Plasma antithrombin concentrations were measured by ELISA (rat Antithrombin III ELISA kit, MyBiosource, San Diego, CA). Plasma prothrombin concentration was measured by gel electrophoresis and immunoblotting, as follows: Equal amounts of PPP were diluted in Laemmli buffer (BIO-RAD, Hercules, CA) with β-mercaptoethanol, resolved on a 10% SDS-polyacrylamide gel (Mini-PROTEAN II, BIO-RAD) and then electrophoretically transferred to a polyvinylidene fluoride membrane (Millipore, Billerica, MA). After blocking with 5% non-fat dry milk solution, membranes were incubated overnight at 1:2000 in primary antibody (Anti-murine Prothrombin, Haematologic Technologies Inc, Essex Junction, VT) followed by corresponding secondary antibody conjugated to horseradish peroxidase. Quantitative determination of protein was performed by autoradiography after revealing the antibody-bound protein by enhanced chemiluminescence reaction (MilliporeSigma, Burlington, MA). The density of the bands on scanned autoradiographs was quantified relative to an identical volume of rat pooled normal plasma using ImageJ (NIH, Bethesda, MD). ELISA and immunoblot antibodies were validated using species-specific positive (purified species-specific protein; Haematologic Technologies, Inc, Essex Juntion, VT) and non-specific protein negative controls. Plasma antithrombin activity was measured using a modified amidolytic method as previously described,^12, 44^ while plasma prothrombin functional activity was determined using a commercially available chromogenic assay (Rox Prothrombin; DiaPharma, West Chester Township, OH).

### Statistical Analyses

The unpaired Student’s t-test was used for single comparisons and one- or two-way ANOVA (analysis of variance) for multiple group comparisons, using SigmaStat software (Systat, San Jose, CA). When a significant difference was identified by ANOVA, post hoc tests were performed using the Student-Newman-Keuls technique. Chi-square or Fisher’s exact test, as appropriate, were used for categorical comparisons. Statistical significance was defined as *P*<0.05. Data are presented as mean ± SE.

## RESULTS

### Both Methylprednisolone and Pioglitazone Alleviate Proteinuria and Hypoalbuminemia

As expected, significant levels of proteinuria (Fig. 1A) and hypoalbuminemia (Fig. 1B) were observed 11 days post-PAN. Treatment with High-MP, Pio, or Pio+Low-MP partially ameliorated PAN-NS. The High-MP and Pio groups had significantly reduced proteinuria compared to untreated PAN rats (*P*<0.05), whereas the Low-MP and Pio+Low-MP groups did not improve. Intriguingly, Pio and High-MP similarly improved proteinuria (73.9% and 69.6% reductions vs. sham, respectively). As expected, plasma albumin levels improved in concert with proteinuria improvement. While Low-MP did not improve hypoalbuminemia, High-MP, Pio, and Pio+Low-MP all improved albumin levels vs. no treatment (*P*<0.05).

**Figure 1:**
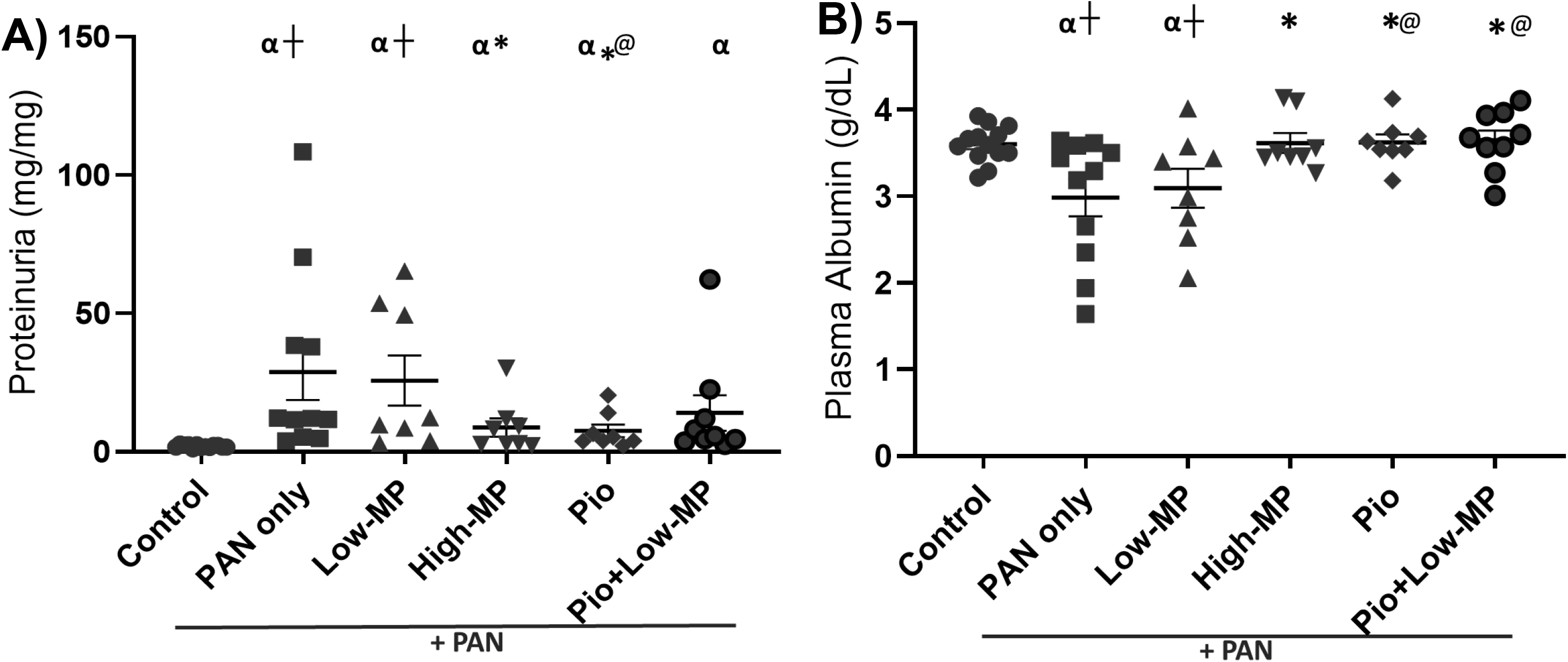
Both Methylprednisolone and Pioglitazone Alleviate Proteinuria and Hypoalbuminemia. Mean +/− SE of **(A)** proteinuria and **(B)** plasma albumin concentration in a PAN-induced rodent model of NS, with/without concomitant methylprednisolone (MP) and/or pioglitazone (Pio) treatment (*n*=8-13/group). α, *, @, †; denotes *P*<0.05 vs Control, PAN only, Low-MP, & High-MP groups, respectively.

### Hypercoagulopathy Improves in Parallel with Nephrosis Following Treatment

As previously demonstrated, proteinuria (*P*=0.005) and hypoalbuminemia (*P*<0.001) were significantly correlated with ETP (Fig. 2).^12^ Successful treatment with either High-MP or Pio reduced ETP to levels similar to control (*P*<0.05 vs. PAN). In contrast, Low-MP and Pio+Low-MP significantly reduced ETP vs. PAN (*P*<0.05), but they did not correct ETP to control values, representing a partial ETP recovery.

**Figure 2:**
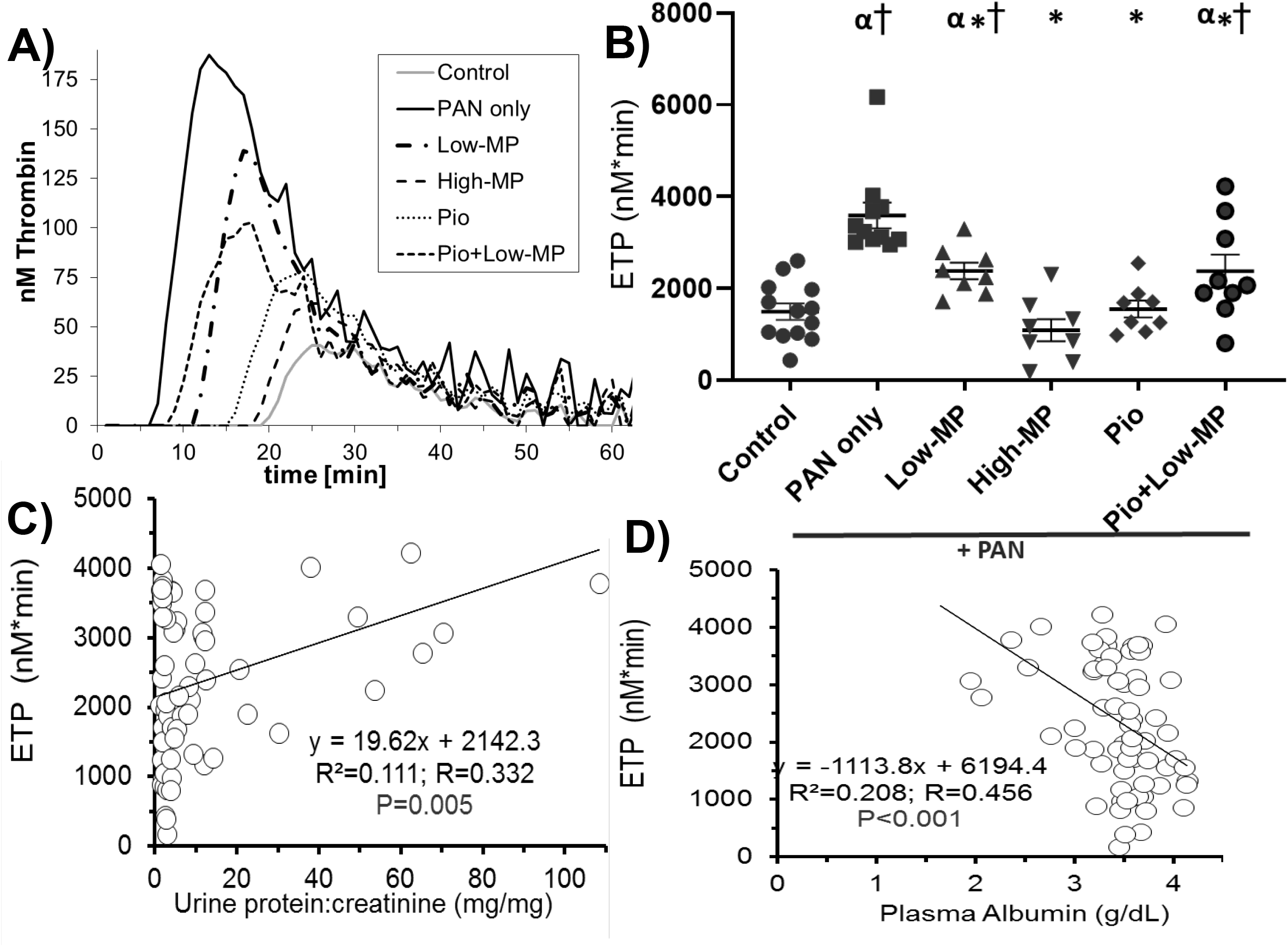
Hypercoagulopathy Improves in Parallel with Nephrosis Following Treatment. **(A)** Representative thrombin generation graphs from one individual rat per group, and graph of mean +/− SE for **(B)** endogenous thrombin potential (ETP), in a PAN-induced rodent model of NS, with/without concomitant methylprednisolone (MP) and/or pioglitazone (Pio) treatment (*n*=8-13/group). (**C-D**) Linear regression analysis correlating disease severity (proteinuria **(C)** and hypoalbuminemia **(D)**) with ETP. α, *, @, †; denotes *P*<0.05 vs Control, PAN only, Low-MP, & High-MP groups, respectively.

### Qualitative Antithrombin Deficit Improves with Treatment Response

We previously demonstrated a qualitative antithrombin deficiency in PAN-NS.^12^ As expected, based on this prior observation, plasma antithrombin antigen (protein) levels were unaffected by either disease or treatment (Fig. 3A-C). In contrast, antithrombin enzymatic activity was modestly improved after treatment with High-MP, Pio, and Pio+Low-MP, such that they were not different from control, but also not significantly higher than in untreated PAN-NS (Fig. 3D). Further, antithrombin activity was significantly correlated with UPC (Fig. 3E; *P*=0.043) but not plasma albumin (Fig. 3F; *P*=0.258). Thus, the groups which showed a response to treatment (improved UPC and plasma albumin) also demonstrated improvements in plasma antithrombin activity. There was no significant correlation between ETP and antithrombin activity or antigen (*P*=0.066 and *P*=0.186, respectively; data not shown). There were also no significant differences in prothrombin antigen (Fig. 4A) or activity (Fig. 4B) levels by treatment group.

**Figure 3:**
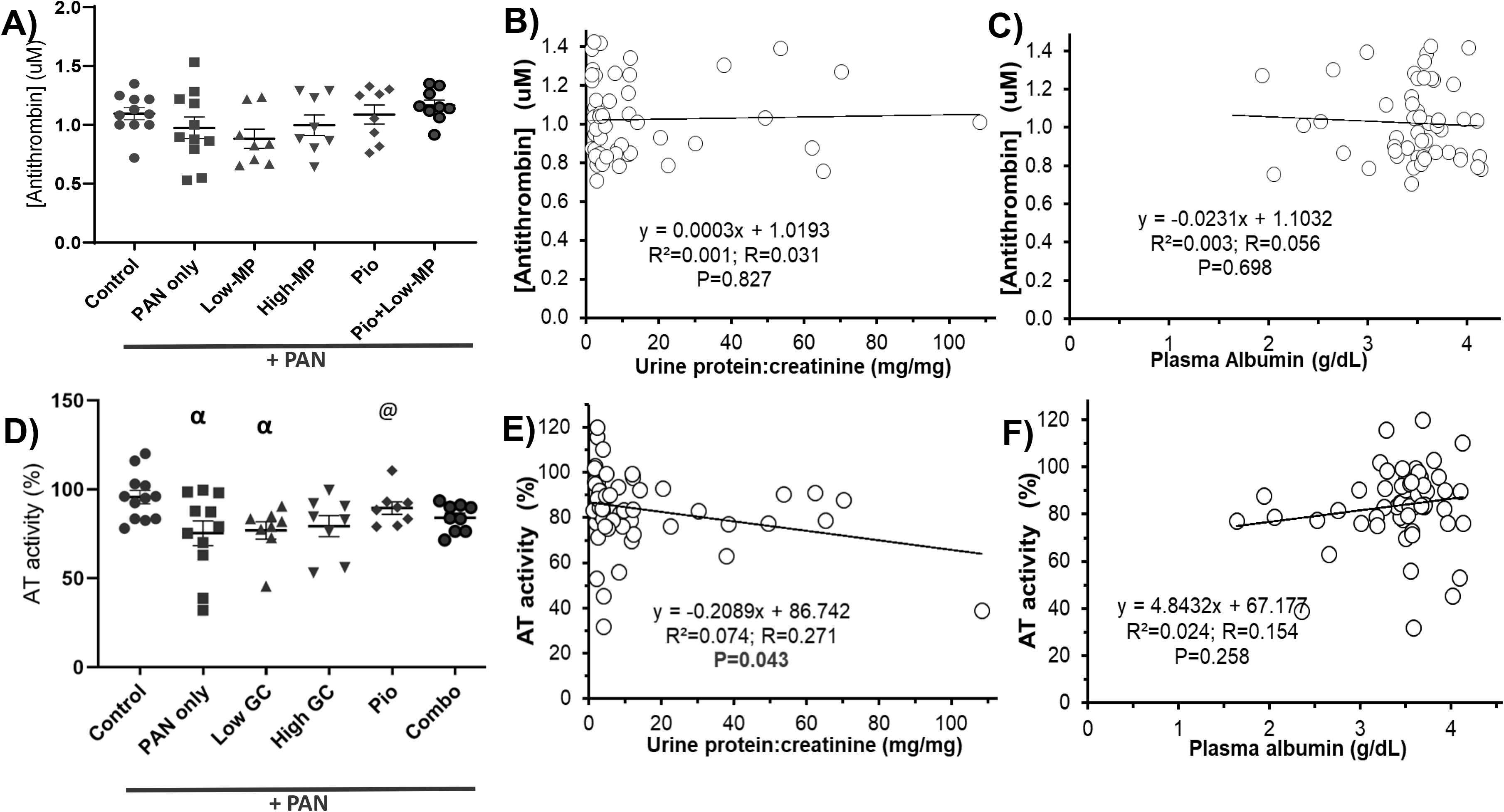
Qualitative Antithrombin Deficit Improves with Treatment Response. Mean +/− SE of plasma AT concentration **(A-C)** and AT activity **(D-F)** in PAN-NS, with/without methylprednisolone (MP) and/or pioglitazone (Pio) treatment, and linear regression analysis with proteinuria **(B, E)** and hypoalbuminemia **(C, F)** (*n*=8-13/group). There was no correlation between ETP and AT activity or AT concentration (*P*=0.066 and *P*=0.186, respectively; data not shown). α, *, @, †; denotes *P*<0.05 vs Control, PAN only, Low-MP, & High-MP groups, respectively.

**Figure 4:**
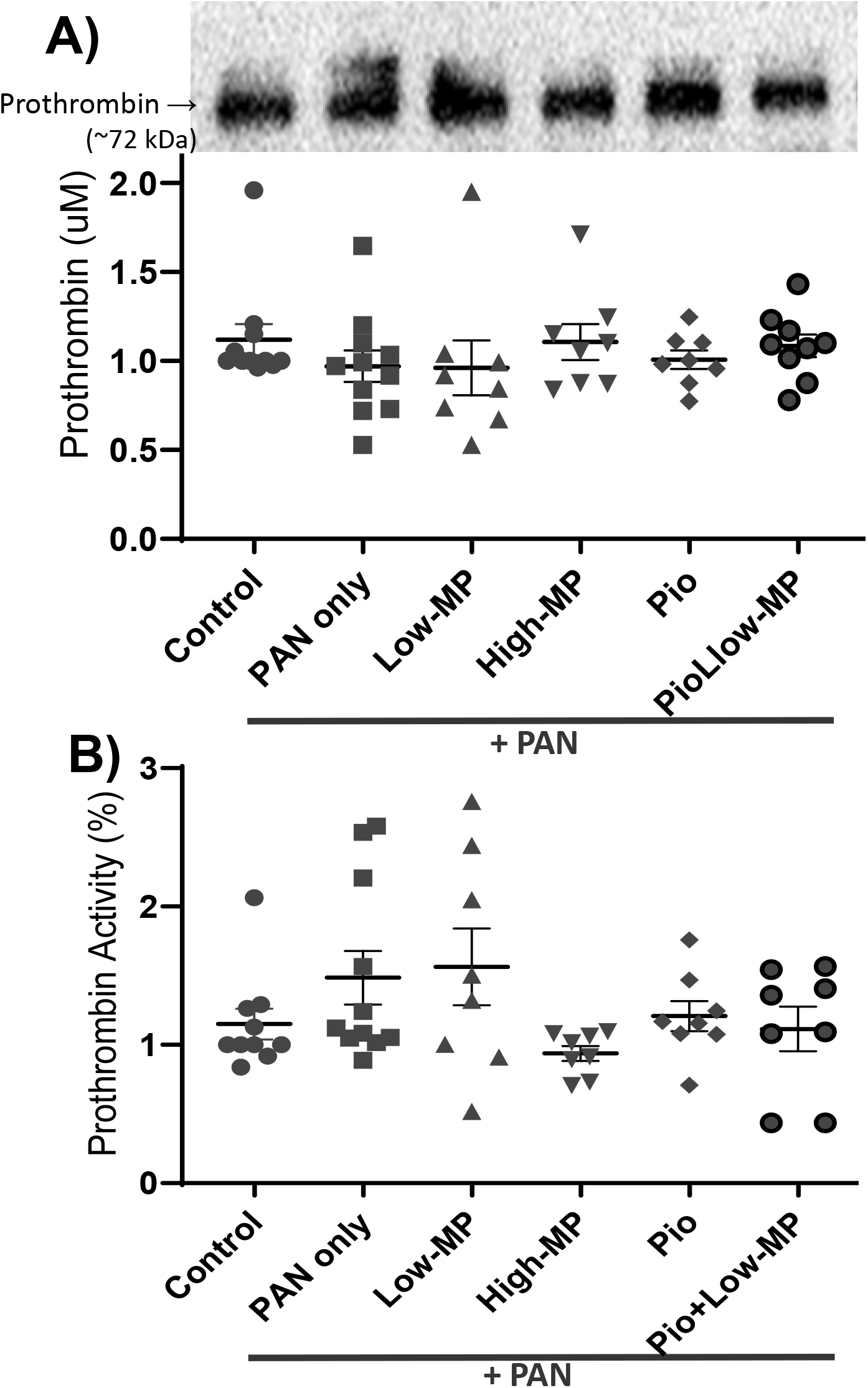
Prothrombin Antigen and Activity are not Altered by PAN-Induced Nephrosis or Treatment. Mean +/− SE of plasma prothrombin concentration (**top panels**) and activity (**bottom panels**) in PAN-induced rodent model of NS, with/without concomitant methylprednisolone (**MP**) and/or pioglitazone (**Pio**) treatment (n=8-13/group).

### Disease and Hypercoagulopathy Responses Persist Across a Broad Range of Disease Severity

Rat PAN-NS severity is variable in a manner dependent on PAN dose and route of administration.^12, 43^ We thus assessed the hypercoagulopathy response across a range of PAN-NS severities. Plasma albumin and UPC were also responsive to High-MP treatment following higher doses of PAN (IV 75 mg/kg or IP 100 mg/kg) (Fig. S1). Similarly, High-MP treatment improved ETP toward control values across the range of tested strategies (Fig. S2). ETP was again significantly correlated with PAN-NS severity. As expected, there were no significant differences in antithrombin antigen levels across the PAN-NS severity spectrum, nor were antigen levels correlated to proteinuria or plasma albumin (Fig. S3). In contrast to antigen levels, antithrombin activity improved with High-MP treatment and was significantly correlated with markers of NS disease severity. Nonetheless, there was no correlation between ETP and antithrombin activity (P=0.592; data not shown).

To further evaluate the hypothesis that NS hypercoagulopathy correlates directly with NS-severity, we performed linear regression analyses on the combined data from all PAN-NS experimental groups presented in this paper (Fig. 5). As expected, in this large combined dataset, proteinuria and hypoalbuminemia were correlated with ETP (*R*^2^=0.211 and 0.139, respectively; *P*<0.001). Plasma antithrombin activity was also significantly correlated to proteinuria and hypoalbuminemia severity (*R*^2^=0.165, *P*<0.001 and *R*^2^=0.110, *P*=0.001; respectively). In contrast to the smaller analyses of each PAN-NS cohort where there was no relationship between antithrombin activity and ETP, in the combined analysis a significant correlation between antithrombin activity and ETP emerged (*R*^2^=0.235, *P*<0.001; Fig. 5E). However, there was still no significant relationship between ETP and antithrombin antigen concentration (*P*=0.309).

**Figure 5:**
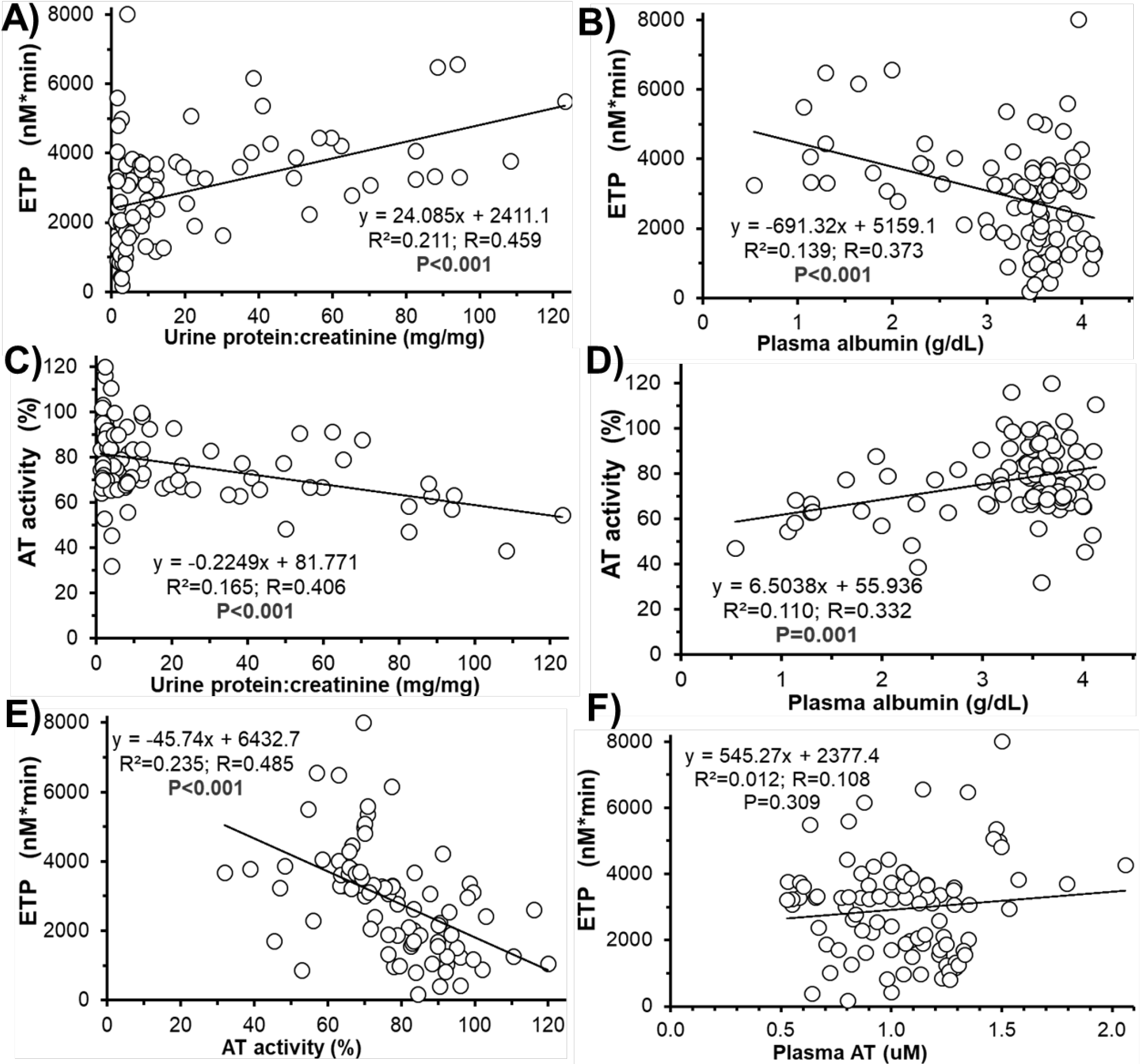
Hypercoagulopathy Responses Persist Across a Broad Range of Disease Severity. Linear regression analysis of disease severity (proteinuria **(A, C)** and hypoalbuminemia **(B, D)**) and coagulation markers (ETP **(A, B)**, AT activity **(C, D)**), in all PAN-NS rats combined (*n*=95). **(E, F)** Linear regression analysis of the relationship between ETP and AT activity **(E)** and AT concentration **(F)** in the combined rat groups (*n*=95).

### Both Methylprednisolone and Pioglitazone Induce Hypercoagulopathy in Non-Nephrotic Rats

As expected, there was no change in proteinuria or plasma albumin, in healthy rats treated with High-MP, Pio, or Pio+Low-MP (Fig. S4). However, all three of these treatments increased ETP in healthy rats (Fig. S5). As expected, in these otherwise healthy animals without PAN-NS, ETP was not correlated with proteinuria or plasma albumin levels (data not shown). Interestingly, however, healthy animals given High-MP also exhibited decreased antithrombin antigen, antithrombin activity, prothrombin antigen, and prothrombin activity (*P*<0.05; Fig. S6). In contrast, Pio and Pio+Low-MP did not cause an antithrombin deficit, and instead led to increased antithrombin protein levels compared to control (*P*<0.05). Antithrombin activity was not correlated to antithrombin protein levels (*P*=0.841) or ETP (*P*=0.118) in these healthy rats. Pio and Pio+Low-MP did not significantly change prothrombin antigen or activity in comparison to healthy control rats.

### Hypercoagulopathy Improves in Children with Steroid-Sensitive, but not Steroid-Resistant Nephrotic Syndrome

Thirty-eight children were enrolled in the PNRC cohort (24 with SSNS and 14 with SRNS; Table S2). As expected, the children with SRNS were significantly older at presentation, otherwise the groups were demographically similar. Proteinuria, plasma albumin, and ETP were not different between the SSNS and SRNS groups at disease presentation (before GC treatment; Fig. 6). As expected, GC therapy significantly improved proteinuria and hypoalbuminemia in children with SSNS (*P*<0.001). ETP was significantly reduced at the followup visit for children with SSNS in comparison to their ETP at presentation (4,505 ± 251 vs. 5,608 ± 298 nM*min, *P*=0.007). In contrast, there was no significant change in ETP for children with SRNS (5,362 ± 377 vs. 5,165 ± 233 nM*min at presentation and follow-up, respectively; *P*=0.661). When combining the disease activity for all these children from both visits, ETP was significantly correlated with both proteinuria (*P*=0.04) and plasma albumin (*P*<0.001) suggesting that hypercoagulopathy is correlated with disease activity in children with NS.

**Figure 6:**
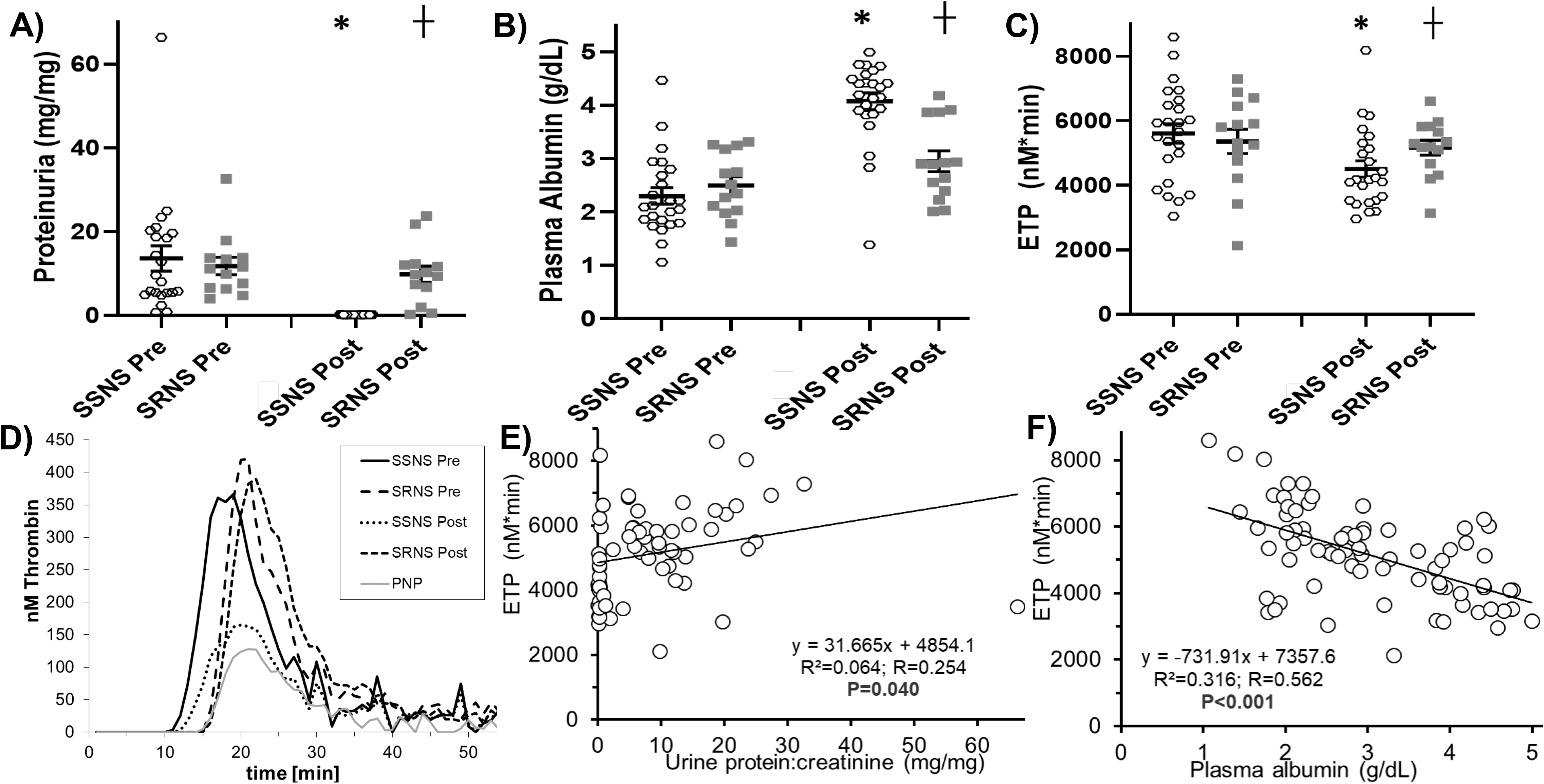
Hypercoagulopathy Improves in Children with Steroid-Sensitive, but not Steroid-Resistant Nephrotic Syndrome. Mean ± SE of proteinuria **(A)**, plasma albumin **(B)**, and ETP **(C)** in childhood steroid-sensitive NS (SSNS; *n*=24) and steroid-resistant NS (SRNS; *n*=14) at disease presentation (“Pre”) and following glucocorticoid treatment (“Post”). **(D)** Representative thrombin generation graphs from one individual per group and a human pooled normal plasma (PNP) control. **(E, F)** Linear regression analysis correlating ETP with proteinuria and hypoalbuminemia, respectively. *, †; denotes *P*<0.05 vs “SSNS Pre”, & “SSNS Post” groups, respectively.

## DISCUSSION

This study investigated the effects of two nuclear receptor agonists, which have disparate mechanisms of action, on endogenous thrombin potential (ETP), a biomarker of venous thromboembolism (VTE) risk in a well-established rodent nephrotic syndrome (NS) model as well as in children with idiopathic NS. Both methylprednisolone (MP), an established frontline childhood NS therapy, and pioglitazone (Pio; an investigational NS therapeutic) significantly reduced proteinuria in rats with puromycin aminonucleoside (PAN) nephrosis and simultaneously improved NS-associated hypercoagulopathy, as measured by ETP. Importantly, Pio was as effective as High-dose MP, suggesting that it may enable a steroid-sparing treatment strategy. These studies further demonstrate that NS disease activity (as determined by proteinuria and hypoalbuminemia) is correlated with ETP, independent of treatment modality. Moreover, in this cohort of children with NS, there was a significant ETP reduction in those with steroid-sensitive NS (SSNS), whereas those with steroid-resistant NS (SRNS) had no ETP improvement from their baseline values. Collectively, these data suggest that treatments which effectively reduce proteinuria may simultaneously reduce NS-associated VTE-risk. Moreover, because ETP is correlated with disease activity, even a partial reduction in proteinuria may provide a secondary clinical benefit by reducing VTE-risk. Finally, this study provides evidence that consideration should be given for inclusion of ETP as a component of composite outcomes when evaluating investigational NS therapeutics in future clinical trials.

These experiments confirm our previous observation that NS-associated hypercoagulopathy is proportional to disease activity and extend this paradigm by showing that hypercoagulopathy improves in parallel with treatment response.^12^ The improvements in ETP were seen using two drugs which act through different members of the nuclear receptor superfamily (MP via glucocorticoid receptor and pioglitazone via PPARγ).^30, 31^ Interestingly, both of these agents enhanced ETP in non-nephrotic rats. Thus, these data strongly suggest that the benefits of these drugs to diminish NS-associated hypercoagulopathy are an indirect consequence of proteinuria (glomerular filtration defect) reduction. Furthermore, these data suggest the possibility that any treatment that effectively reduces NS severity may also indirectly ameliorate NS-associated hypercoagulopathy. This concept is further supported by the human data showing that children with SSNS had improved ETP from their baseline values, whereas children with SRNS did not. Thus, an indirect benefit of effective NS therapy is expected to be reversal of hypercoagulopathy and thus, decreased VTE-risk. These data are consistent with previous studies demonstrating improvement of both procoagulant and anticoagulant protein levels and enzymatic activities between active NS and NS in remission.^45–48^

NS complications may occur due to the disease itself or as a consequence of adverse therapeutic events. Glucocorticoids (i.e. MP and prednisolone) remain the most common frontline therapy for idiopathic childhood NS, but are associated with well-recognized and potentially serious side effects.^29^ In addition, ~10-15% of childhood NS cases are steroid-resistant (SRNS) and some data suggests that the prevalence of SRNS is rising.^49^ Moreover, NS continues to be a leading cause of ESKD despite GC and alternative immunosuppressant therapies.^24–26, 28, 50^ Thus, a critical need remains to develop more targeted and less toxic therapies to improve NS outcomes. In this regard, we previously demonstrated that Pio reduced proteinuria in PAN-induced NS (both independently and in combination with reduced-dose MP), suggesting that Pio may enable a steroid-sparing NS treatment strategy.^30^ The present study confirms and extends these observations to additional PAN-NS groups and demonstrates that Pio treatment (alone or in combination with low-dose MP) results in a partial proteinuria reduction of (73.9% or 51.4%, respectively) that was similar to conventional, high-dose MP (69.6%). In contrast, low-dose MP without Pio was not beneficial. Furthermore, this study is the first to describe the effects of Pio treatment on hemostasis in NS, demonstrating that effective proteinuria reduction corelated with a complete alleviation of NS-associated hypercoagulopathy, as evidenced by ETP correction to values indistinguishable from those of control rats. Thus, Pio (alone or in combination) successfully reduced both proteinuria and NS-associated hypercoagulopathy. Therefore, Pio may have significant potential as a novel NS therapy, to simultaneously reduce disease severity (primary outcome) and limit both steroid-related side effects and VTE-risk (secondary outcomes). Pio may be particularly beneficial for childhood NS, where there is less risk of thiazolidinedione-mediated heart failure.^51–53^ Thus, a multicenter randomized controlled trial assessing the clinical benefits of Pio in the treatment of childhood idiopathic NS should include ETP as a component of the composite outcome.

It has long been suggested that GC administration induces procoagulant effects in otherwise healthy individuals.^36^ Although studies in healthy volunteers are appropriate for detecting unconfounded effects of GC, in clinical practice GC are most commonly used for inflammatory diseases and in surgical settings, and thus most published literature on the effects of MP on hemostasis are confounded by the simultaneous effects of the underlying disease processes.^34, 38, 54, 55^ Nonetheless, there appears to be little doubt that patients treated with GC have higher VTE-risk. For example, a recent, large population-based case-control study found that systemic GC therapy (including MP) was associated with increased VTE-risk, an observation that persisted after adjustment for underlying disease severity, for both inflammatory and non-inflammatory conditions.^55^ Results from previous studies suggest that alterations in coagulation balance are dependent on both GC drug and dose. A variety of hemostatic effects have been detected including: increased platelet aggregation, shortened partial thromboplastin times, increased levels of factors V, VII, VIII, XI, and fibrinogen, which are associated with enhanced arterial thrombosis *in vivo*.^34–38^ However, data regarding the effects of GC on overall hemostatic balance, as determined by ETP and other global coagulation assays are lacking. Only one previous study employed a thrombin generation assay in healthy adults administered prednisolone (0.5 mg/kg/day orally) for 10 days.^35^ That investigation revealed increases in peak thrombin and velocity-index, but not ETP. However, there were unexpectedly high thrombin generation parameters in the placebo group (at baseline) that limited the interpretation of the data. In the present study, healthy rats treated with MP were hypercoagulable by ETP. In addition, both qualitative and quantitative AT defects were present in healthy rats treated with High-MP, but not with Pio or Pio+Low-MP therapy. Taken together the present and previously published data suggest that GCs induce a procoagulant state in healthy subjects.

In contrast to the procoagulant effects of GC, thiazolidinediones may have anticoagulant effects that include decreased factor VII, plasminogen activator inhibitor-1, von Willebrand factor, and platelet activation.^39–42^ In healthy rats administered low (1 mg/kg) or high (10 mg/kg) dose Pio for 10 days, decreased platelet aggregation and delayed arterial thrombus formation were observed.^56^ These beneficial effects were likely secondary to increases in aortic wall expression of thrombomodulin and constitutively-expressed nitric oxide synthase (cNOS). In contrast to these anticoagulant effects, the present data demonstrate that both Pio and Pio+Low-MP increase thrombin generation (ETP) when administered to healthy animals. However, *in vivo* thrombosis modeling suggests that PPARγ signaling is antithrombotic, suggesting that the antithrombotic effects of Pio on the vessel wall dominate the prothrombotic ETP elevations found in the plasma compartment.^57, 58^ Consistent with this interpretation, the clinical evidence suggests that Pio has an overall positive safety profile and lack of VTE-risk.^59, 60^

Although it has been suggested that GC therapy may contribute to increased VTE-risk during NS, results from epidemiologic studies have generally failed to support a clear link.^5, 54, 61, 62^ Whereas SRNS patients are known to have a higher VTE-risk than those with SSNS,^54, 63^ the factors contributing to this discrepancy are not yet fully elucidated. However, the present data demonstrating that the acquired NS-hypercoagulopathy is highly correlated with disease activity, during both disease and treatment, suggest that the persistent proteinuria associated with SRNS corresponds with persistent hypercoagulopathy. Indeed, the ETP data from this small cohort of children with SSNS vs. SRNS supports this hypothesis, but should be extended to a larger patient cohort before reaching definitive conclusions. If confirmed, this paradigm would further support the idea that patients with persistent NS should remain on anticoagulant prophylaxis whereas it may be safe to discontinue anticoagulation in patients who have achieved a sustained complete remission.

We previously demonstrated a qualitative antithrombin deficiency in PAN-NS.^12^ In the present study, antithrombin activity improved in concert with treatment response, improving to healthy control levels in animals that responded to therapy. Interestingly, there was no direct correlation between antithrombin activity and ETP in the individual rat experiments (*n*=58 & 38). Only when combining all of the rat groups together (*n*=95) was a correlation between ETP and antithrombin activity uncovered, suggesting that in PAN-NS, the qualitative antithrombin deficit likely makes only a minor contribution to increased thrombin generation. This interpretation is also consistent with the appearance of the thrombin generation curves wherein the ETP increases appear to originate from shortened lag-times and increased peak thrombin values, rather than the tail prolongation that has been demonstrated in antithrombin deficient plasma.^64^

In these experiments, variations of the single-dose PAN-NS model were used to investigate treatment effects at peak disease severity (day 11), therefore Pio and/or MP treatment was commenced the same day as PAN administration. Future studies investigating the efficacy of Pio and/or MP to reduce established PAN-NS as well as other NS models are thus warranted to further elucidate the potential benefits of these treatments on NS-associated hypercoagulopathy. Nonetheless, the data demonstrating improved ETP with high-dose MP across a wide range of proteinuria severity suggests that the relationship between disease reduction and ETP improvement will be generalizable to other models. Moreover, the proteinuria levels induced in the Pio experiments are on par with those we previously demonstrated to exacerbate thrombosis in PAN-NS,^12^ therefore the Pio-induced reductions in ETP are likely to be physiologically relevant.

Both pioglitazone and methylprednisolone simultaneously improved proteinuria and NS-associated hypercoagulopathy. Pioglitazone enabled a steroid-sparing treatment strategy in the PAN-NS rat model. These data confirm our previous observation that NS-hypercoagulopathy is proportional to disease severity and, importantly, extend these findings to show that hypercoagulopathy improves in concert with therapeutic response in NS. Even a partial disease response (i.e. partial NS remission) may thus reduce hypercoagulopathy and diminish clinical VTE-risk. Children with steroid resistant NS are persistently hypercoagulopathic and may thus benefit from prolonged anticoagulation. Because these data suggest that any treatment modality that effectively reduces NS disease activity may ameliorate NS-associated hypercoagulopathy, hemostatic assays (e.g. ETP) should be included as valuable secondary outcome variables for use in NS clinical trials employing composite outcomes.

## APPENDIX

Members of the Pediatric Nephrology Research Consortium (PNRC; formerly the Midwest Pediatric Nephrology Consortium)

Nationwide Children’s Hospital and The Ohio State University, Columbus, OH: J Mahan, H Patel, RF Ransom

Medical College of Wisconsin, Milwaukee, WI: C Pan

The Hospital for Sick Children, Toronto, ON: DF Geary

West Virginia University, Charleston, WV: ML Chang

University of North Carolina, Chapel Hill, NC: KL Gibson

Louisiana State University, New Orleans, LA: FM Iorember

University of Iowa Stead Family Children’s Hospital, Iowa City, IA: PD Brophy

Children’s Mercy Hospital, Kansas City, MO: T Srivastava

Emory University School of Medicine, Atlanta, GA: LA Greenbaum

## Author Contributions

APW, KJW, JK, and MAC conducted experiments, analyzed data, prepared figures, and wrote the manuscript; SA analyzed data, prepared figures, and edited the paper; WES and BAK provided animals and reagents, edited the paper, and were responsible for overseeing and coordinating the study. The PNRC authors (Appendix) enrolled, treated, and contributed the patient data without which this study would have been impossible.

## Acknowledgments

The authors thank the patients who contributed blood samples through the PNRC studies. Portions of this work were previously presented at the 12^th^ International Podocyte Conference, Montreal, Canada (May 2018) and the American Society of Nephrology Kidney Week, San Diego, CA (October 2018).

SA is supported by an American Heart Association Career Development Award (CDA34110287). This work was supported by National Institutes of Health National Institute of Diabetes and Digestive and Kidney Diseases grants K08DK103982 (BAK), and R01DK095059 (WES); and the George & Elizabeth Kelly Foundation (Lewis Center, OH; BAK). The content is solely the responsibility of the authors and does not necessarily represent the official views of the National Institutes of Health.

## Disclosures

The authors declare no competing financial interests.

## Supplemental Material

Supplementary Figures 1 – 6

Supplementary Tables 1 – 2

## Supplementary Figures

**Figure S1:**
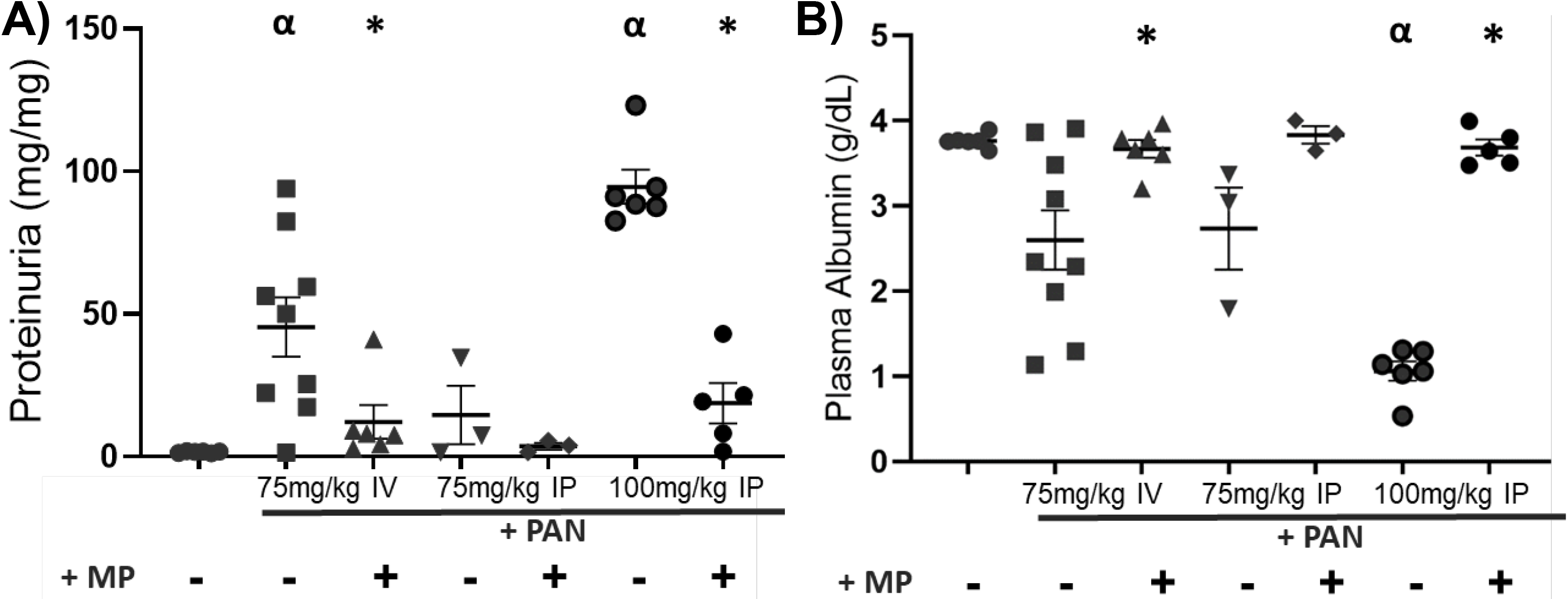
PAN Nephrotic Rats Are Responsive to High-Dose Methylprednisolone Treatment. Mean +/− SE of **(A)** proteinuria and **(B)** plasma albumin concentration in a PAN-induced nephrotic rats (*n*=4-8/group). Varying disease severity was induced in male Wistar rats by a single intravenous (IV) or intraperitoneal (IP) injection of 75 or 100 mg/kg PAN. Steroid sensitivity was confirmed after 11 days of treatment with either high-dose methylprednisolone (+MP; 15 mg/kg) or sham saline (-). α, *, @; denotes *P*<0.05 vs Control & PAN alone, respectively.

**Figure S2:**
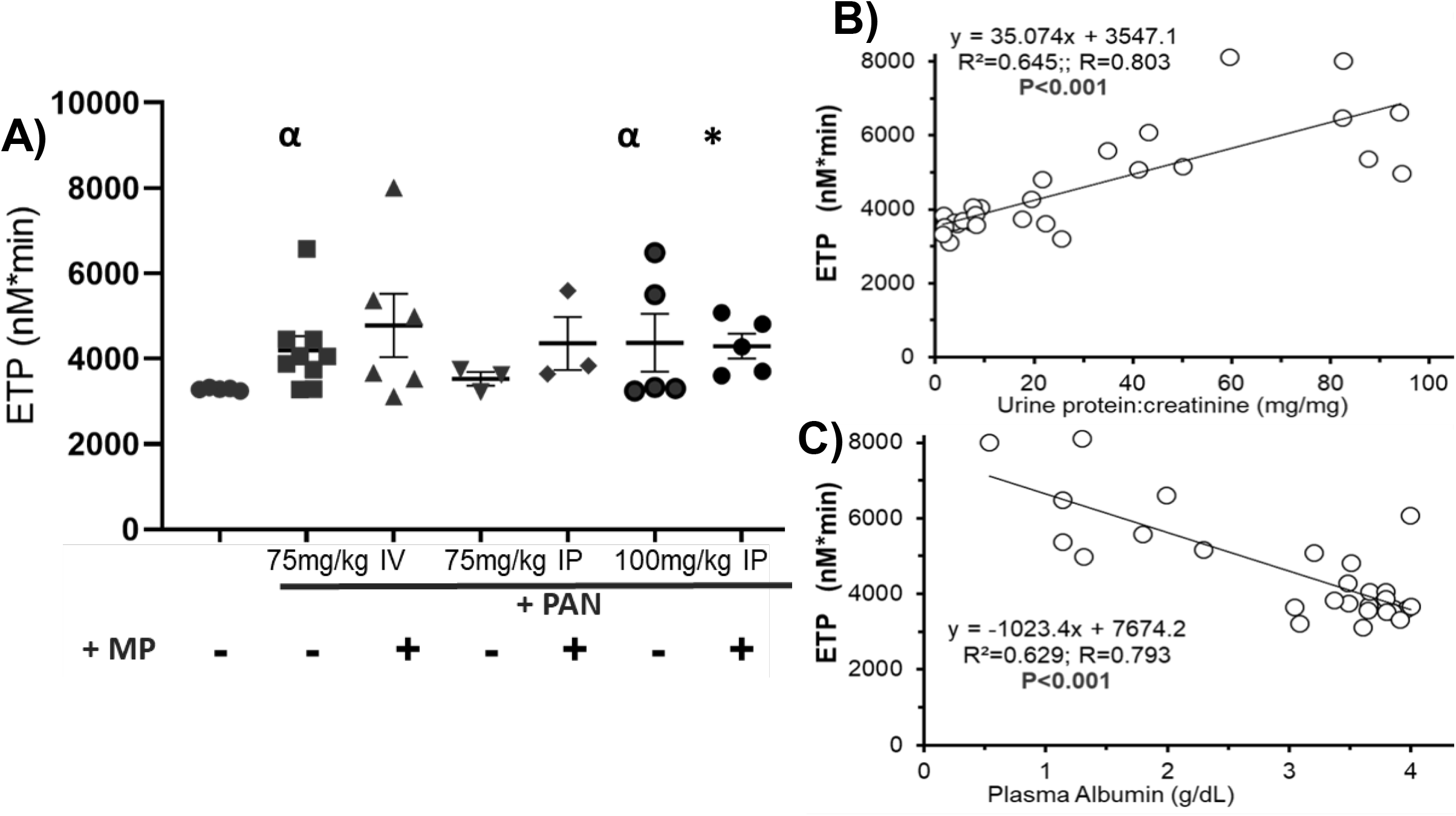
High-Dose Methylprednisolone Corrects Thrombin Generation in Nephrotic Syndrome Over a Wide Range of PAN-Induced Disease Severity. Mean +/− SE of Endogenous Thrombin Potential **(A)** in rats made nephrotic with a single intravenous (IV) or intraperitoneal (IP) injection of 75 or 100 mg/kg PAN injection, and then treated with high-dose methylprednisolone (+MP; 15 mg/kg) or sham saline (-) for 11 days (*n*=4-8/group). ETP was significantly correlated with proteinuria and hypoalbuminemia **(B, C)**. α, *; denotes P<0.05 vs Control & PAN alone, respectively.

**Figure S3:**
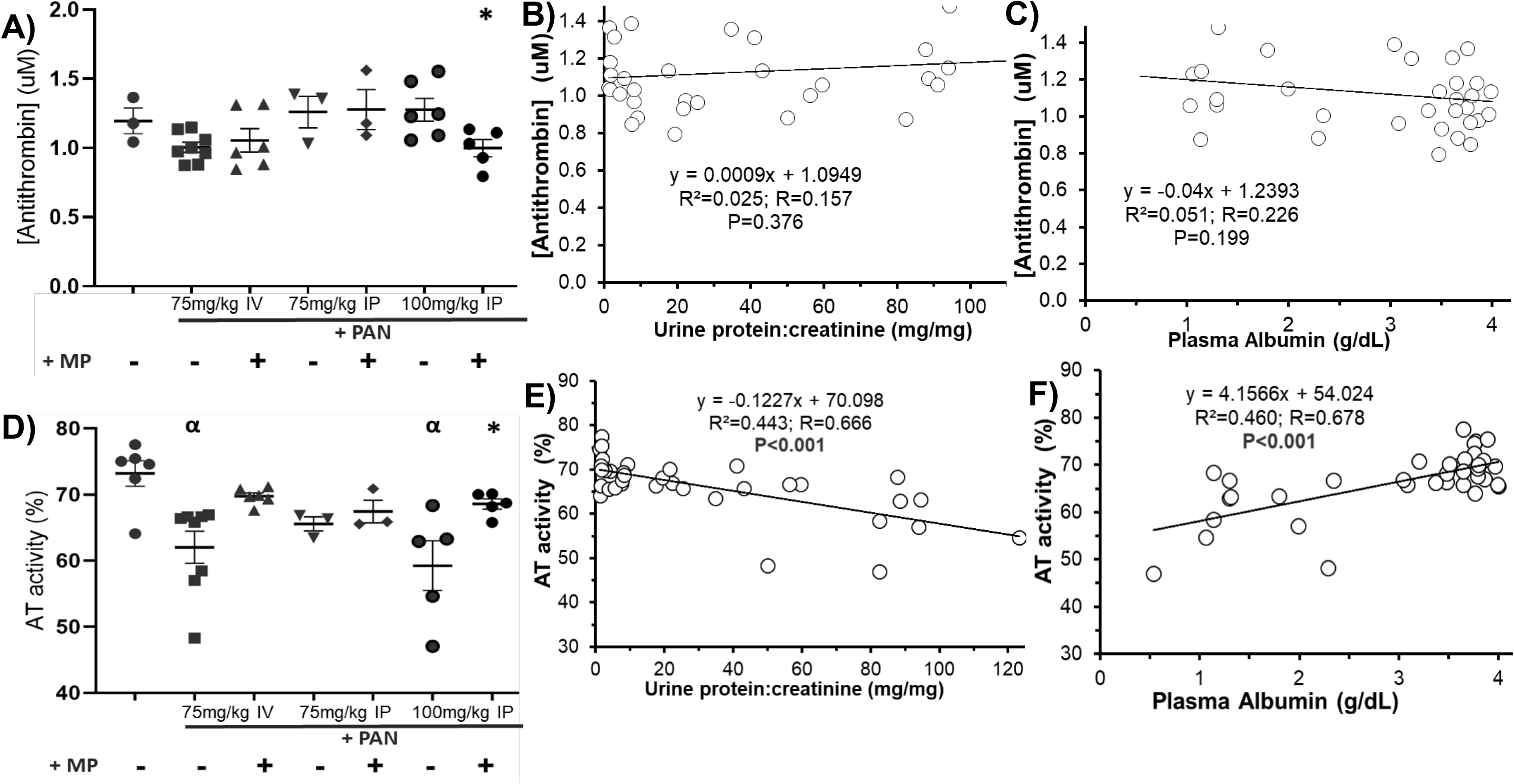
Antithrombin Deficit Improves with High Dose Methylprednisolone Treatment Response, Over A Wide Range of PAN-Induced Disease Severity. Mean +/− SE of plasma antithrombin (AT) antigen **(A-C)** & AT activity **(D-F)** in a PAN-induced rodent model of NS treated with high-dose methylprednisolone (+MP; 15 mg/kg) or sham saline (-) for 11 days (*n*=4-8/group). Plasma AT antigen and activity were significantly correlated with disease severity. α, *; denotes P<0.05 vs Control & PAN alone, respectively.

**Figure S4:**
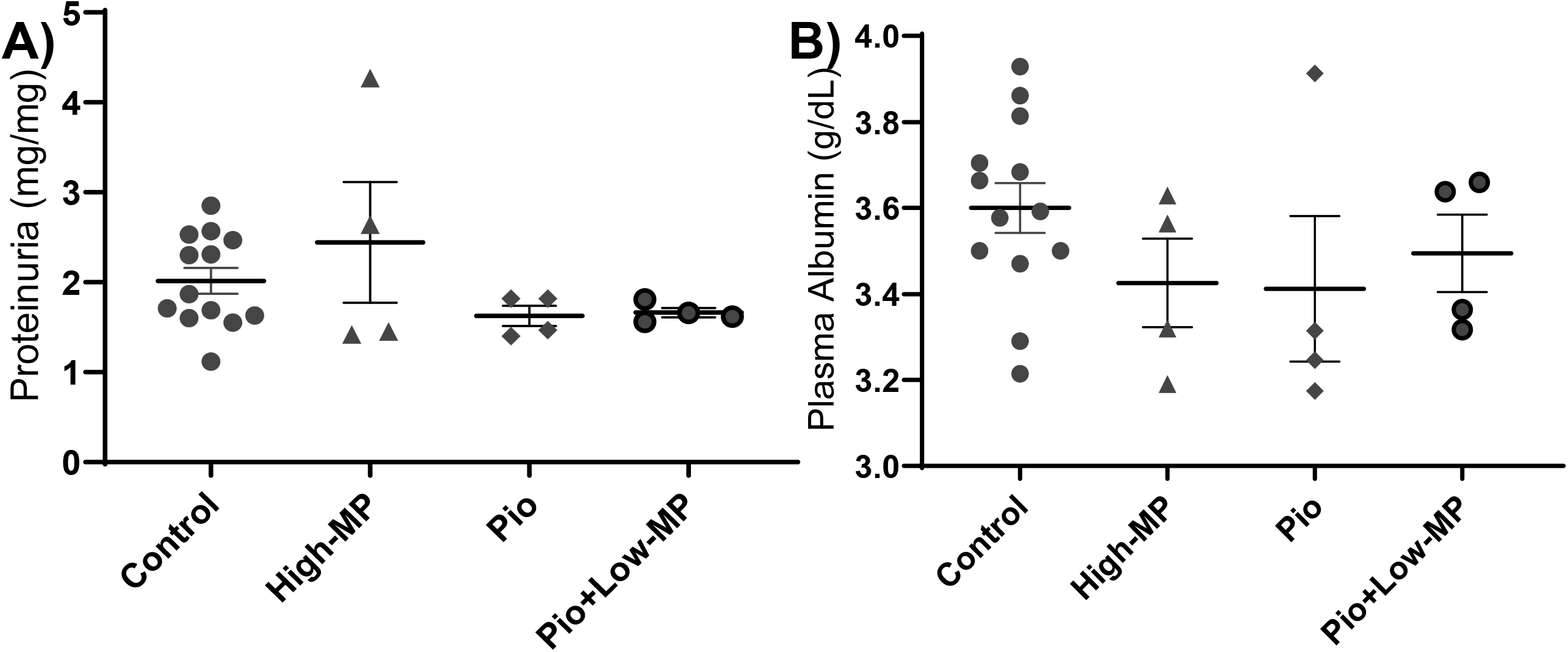
No Differences in Proteinuria or Plasma Albumin of Healthy Rats Treated with Methylprednisolone and Pioglitazone Alone or in Combination with Methylprednisolone. Mean +/− SE of **(A)** proteinuria and **(B)** plasma albumin concentration in healthy Wistar rats treated with saline (Control), high-dose methylprednisolone (MP; 15mg/kg), pioglitazone (Pio), or combination therapy (Pio+Low-GC) (*n*=4-13/group).

**Figure S5:**
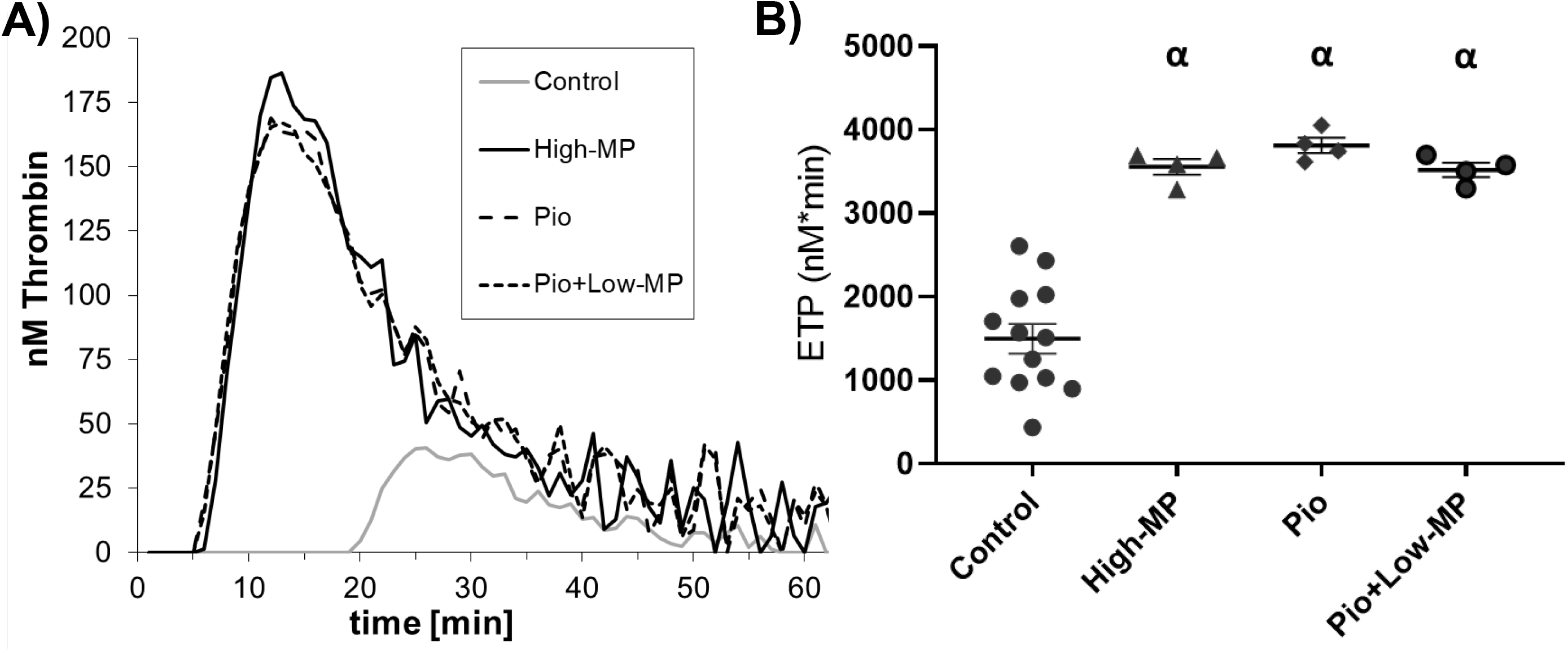
Healthy Rats Treated with Methylprednisolone and Pioglitazone Alone or in Combination Exhibit Elevated Endogenous Thrombin Potential. **(A)** Representative thrombin generation graphs from one individual rat per group, & graph of mean +/− SE of **(B)** Endogenous Thrombin Potential, in healthy Wistar rats treated with saline (Control), high-dose methylprednisolone (MP; 15mg/kg), pioglitazone (Pio), or combination therapy (Pio+Low-GC) (*n*=4-13/group). α; denotes *P*<0.05 vs Control.

**Figure S6:**
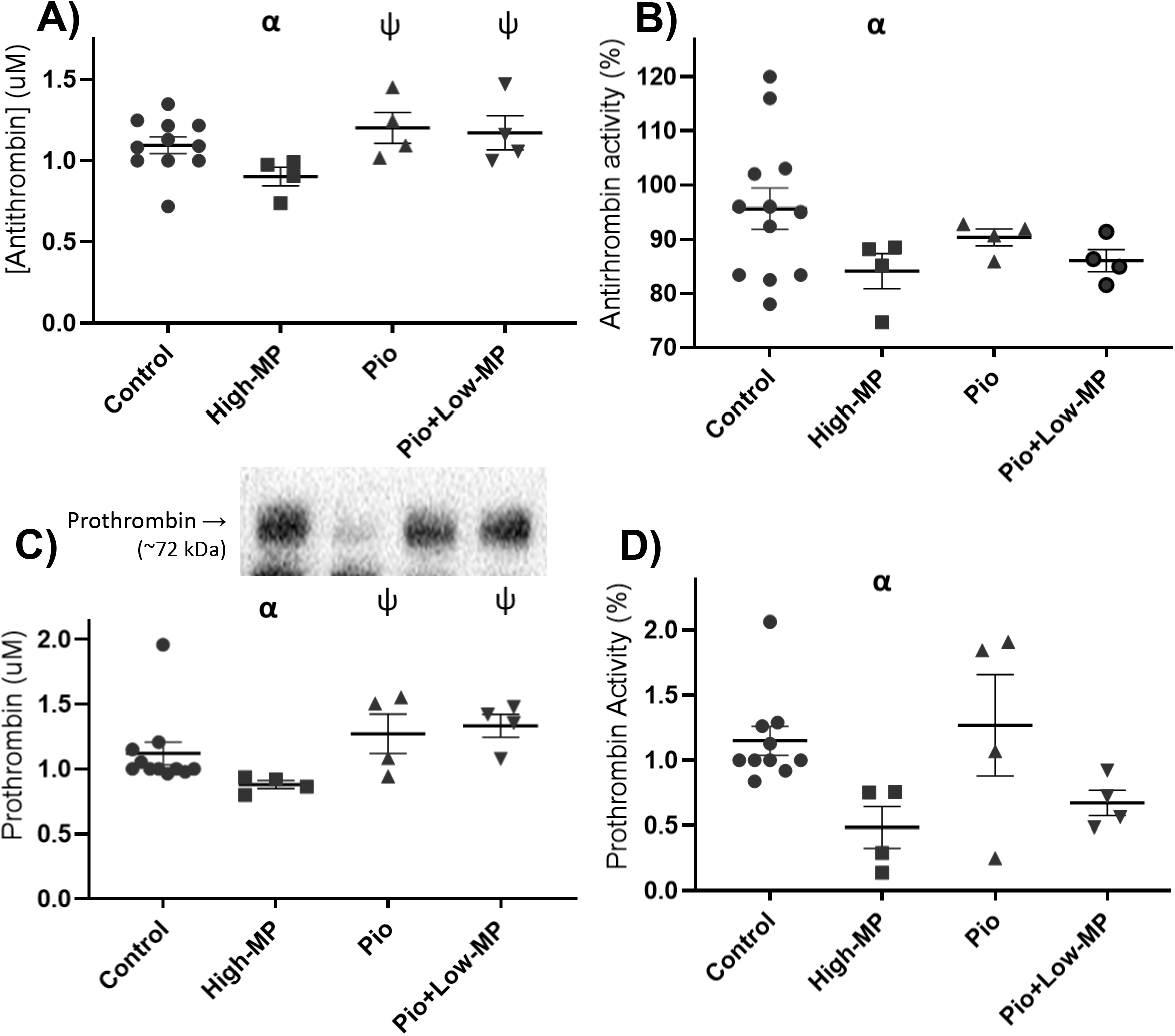
Antithrombin Antigen and Activity Defect and Hypoprothrombinemia in Healthy Rats Treated with High Dose Methylprednisolone. Mean +/− SE of antithrombin **(A)** or prothrombin **(C)** <u>antigen</u> levels, or antithrombin **(B)** or prothrombin **(D)** enzymatic <u>activities</u> in plasma from healthy Wistar rats treated with saline (Control), high-dose methylprednisolone (MP; 15mg/kg), pioglitazone (Pio), or combination therapy (Pio+Low-GC) (n=4-13/group). α, †; denotes P<0.05 vs Control & High-MP groups, respectively.

## Supplementary Tables

**Table S1:**
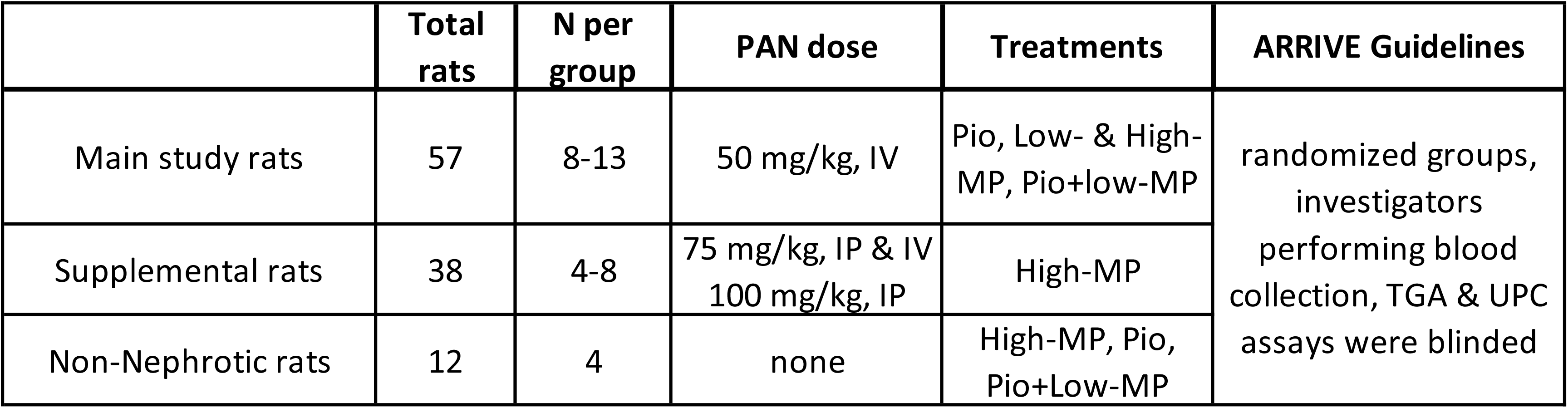
ARRIVE Guidelines for Rat Experiments. IV: intravenous tail vein injection; IP: intraperitoneal injection; Low- and High-MP: 5 and 15 mg/kg methylprednisolone, respectively; Pio: 10 mg/kg pioglitazone via oral gavage; ARRIVE: Animal Research: Reporting *In Vivo* Experiments

**Table S2:**
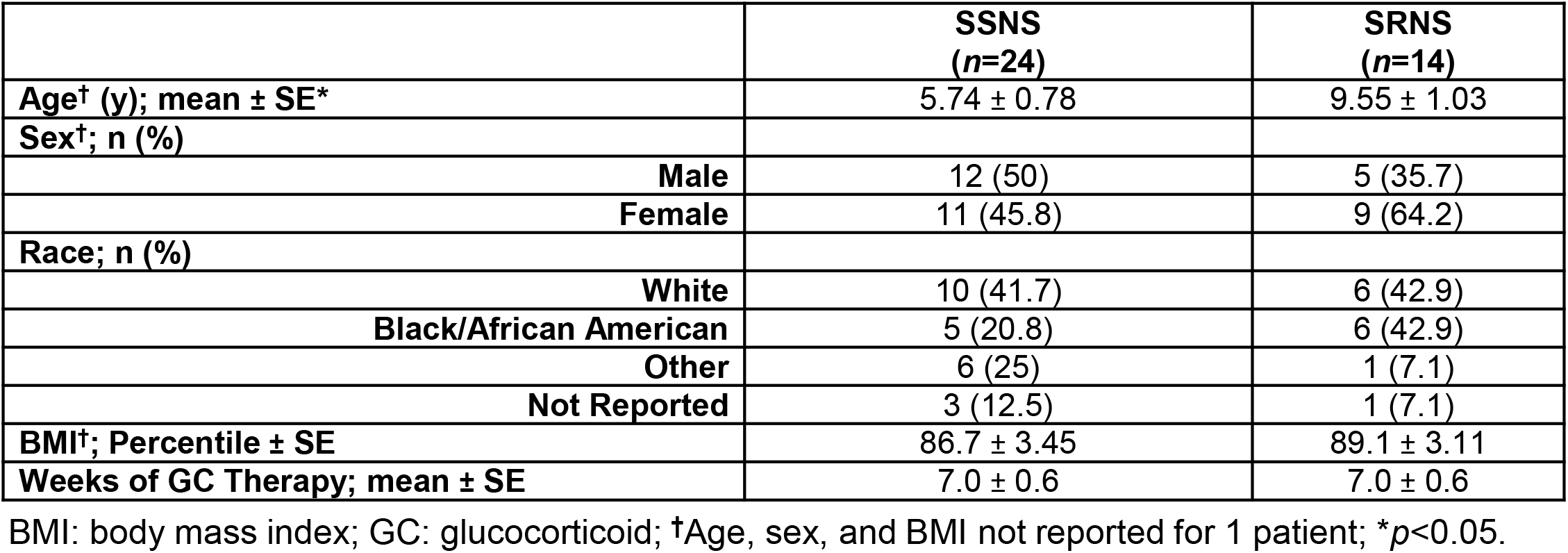
Pediatric Nephrology Research Consortium Cohort Demographics. BMI: body mass index; GC: glucocorticoid; †Age, sex, and BMI not reported for 1 patient; \**p*<0.05.

|  | **SSNS**<br>**(*n*=24)** | **SRNS**<br>**(*n*=14)** |
| --- | --- | --- |
| **Age<sup>†</sup> (y); mean ± SE\*** | 5.74 ± 0.78 | 9.55 ± 1.03 |
| **Sex<sup>†</sup>; n (%)** |  |  |
| **Male** | 12 (50) | 5 (35.7) |
| **Female** | 11 (45.8) | 9 (64.2) |
| **Race; n (%)** |  |  |
| **White** | 10 (41.7) | 6 (42.9) |
| **Black/African American** | 5 (20.8) | 6 (42.9) |
| **Other** | 6 (25) | 1 (7.1) |
| **Not Reported** | 3 (12.5) | 1 (7.1) |
| **BMI<sup>†</sup>; Percentile ± SE** | 86.7 ± 3.45 | 89.1 ± 3.11 |
| **Weeks of GC Therapy; mean ± SE** | 7.0 ± 0.6 | 7.0 ± 0.6 |

